# Genomic architecture of codfishes featured by expansions of innate immune genes and short tandem repeats

**DOI:** 10.1101/163949

**Authors:** Ole K. Tørresen, Marine S. O. Brieuc, Monica H. Solbakken, Elin Sørhus, Alexander J. Nederbragt, Kjetill S. Jakobsen, Sonnich Meier, Rolf B. Edvardsen, Sissel Jentoft

## Abstract

**Background:** Increased availability of genome assemblies for non-model organisms has resulted in invaluable biological and genomic insight into numerous vertebrates including teleosts. The sequencing and assembly of the Atlantic cod (*Gadus morhua*) genome and the genomes of many of its relatives (Gadiformes) demonstrated a shared loss 100 million years ago of the major histocompatibility complex (MHC) II genes. The recent publication of an improved version of the Atlantic cod genome assembly reported an extreme density of tandem repeats compared to other vertebrate genome assemblies. Highly contiguous genome assemblies are needed to further investigate the unusual immune system of the Gadiformes, and the high density of tandem repeats in this group.

**Results:** Here, we have sequenced and assembled the genome of haddock (*Melanogrammus aeglefinus)* - a relative of Atlantic cod - using a combination of PacBio and Illumina reads. Comparative analyses uncover that the haddock genome contains an even higher density of tandem repeats outside and within protein coding sequences than Atlantic cod. Further, both species show an elevated number of tandem repeats in genes mainly involved in signal transduction compared to other teleosts. An in-depth characterization of the immune gene repertoire demonstrates a substantial expansion of *MCHI* in Atlantic cod compared to haddock. In contrast, the Toll-like receptors show a similar pattern of gene losses and expansions. For another gene family associated with the innate immune system, the NOD-like receptors (NLRs), we find a large expansion common to all teleosts, with possible lineage-specific expansions in zebrafish, stickleback and the codfishes.

**Conclusions:** The generation of a highly contiguous genome assembly of haddock revealed that the high density of short tandem repeats as well as expanded immune gene families is not unique to Atlantic cod – but most likely a feature common to all codfishes. A shared expansion of *NLR* genes in teleosts suggests that the *NLRs* have a more substantial role in the innate immunity of teleosts than other vertebrates. Moreover, we find that high copy number genes combined with variable genome assembly qualities may impede complete characterization, i.e. the number of *NLRs* might be underestimates in the different teleost species.

## Background

Recent advances of state-of-the-art genomic tools have resulted in a multitude of whole genome sequencing projects targeting non-model organisms. This has created a new understanding of the genomic underpinnings of the biology of these species and their adaptation to the environment [1]. Examples include the adaptive radiation of African cichlids [2], adaptation to salinity in European sea bass and Atlantic herring [3,4] and drastic morphological changes in pipefish and seahorses [5,6]

The species-rich order Gadiformes, i.e. codfishes and related species, comprises some of the most commercially important fish in the world such as Alaska pollock (*Gadus chalcogrammus*), Atlantic cod (*Gadus morhua*), saithe (*Pollachius virens*) and haddock (*Melanogrammus aeglefinus)* [7,8]. Recent reports have shown that this lineage has undergone dramatic evolutionary changes within its immune system compared to other vertebrates, with a loss of the major histocompatibility complex (MHC) II genes, in the lineage leading to the Gadiformes 105-85 million years ago [9,10]. Additionally, other immune related genes have likely been lost prior to this event, i.e. the Toll-like receptor (*TLR*) *5* 151-147 million years ago and the Myxovirus resistance gene (*Mx*) 126-104 million years ago [11]. A detailed characterization of the TLR gene repertoire – belonging to the pattern recognition receptors (PRRs) family and an important component of the innate immunity [12] – within the Gadiformes lineage revealed specific losses and several expansions [10,13]. Some of these lineage-specific expansions, i.e. TLR8, TLR22, TLR25 and in particular TLR9, were further correlated to the loss of *MHCII* and species latitudinal distributions [14]. An extreme expansion of *MHCI* genes – with more than 100 copies in some species – is another peculiarity of the immune system that Atlantic cod shares with many of the other gadiform species [9]. It has been suggested that some of these *MHCI* genes have taken on a more *MHCII*-like function through cross-presentation; i.e. compensating for the loss of the *MHCII* genes [15]. Taken together, these discoveries suggest that the loss of *MHCII* has fostered immunological innovation - through the altered *TLR* and *MHCI* gene repertoire – within the Gadiformes order.

Another important PRR family is the NOD-like receptors (NLR) class of proteins (also called NACHT- domain- and leucine-rich-repeat-containing receptors or nucleotide-binding domain and leucine-rich-repeat-containing receptors). These recognise microbial products and danger-associated molecular patterns [16]. The NLRs are a large class of intracellular immune receptors in animals [17]. Many species with a classic adaptive immune system contain relatively few *NLR* genes (around 20-30), such as mammals [16,18]. Species without an adaptive immune system, such as cnidarians [19] and the purple sea urchin [20], contain large numbers of *NLRs* (up to 300). Investigations into the *NLRs* repertoire of teleosts indicate different numbers of *NLRs* in different species, e.g. a possible lineage-specific expansion in zebrafish [18].

The major impediment for creating highly contiguous genome assemblies is repeated sequences [21]. For assemblies created solely from short Illumina reads (100-250 bp compared to 800-900 bp for Sanger) these repeated sequences could lead to fragmented assemblies missing important information, such as particular exons or whole genes [22]. With long-read sequencing (10,000 bp and longer as provided by PacBio and Oxford Nanopore), most of the repeats would be spanned, and highly contiguous assemblies surpassing the earlier Sanger based assemblies in quality are possible [23-25]. Highly contiguous assemblies are a prerequisite for in-depth characterization and comparative studies of complex and multi-copy immune gene families (see [13]). Recently, a new version of the Atlantic cod genome assembly was generated by a combination of long read and conventional short read technologies, with substantial contiguity improvements compared to the previous version [26]. The improved assembly revealed an unusual high 10^-8,^ to 10^-2,^ mutations per locus per generation [28], and are density of short tandem repeats (STRs, DNA motifs of 1–10 bp repeated in tandem) compared to other vertebrates [26]. STRs mutate at high rates [27], in humans from 10-located in about 4,500 human genes [29]. Expression of about 2,000 human genes is significantly associated with STR length variation in regulatory regions [30]. The Atlantic cod has about three times the density and frequency of STRs compared to humans, both in coding and non-coding regions [26]. Notably, this suggests that a substantial higher fraction of genes are associated with STRs in Atlantic cod compared to the human genome. These STRs might facilitate evolvability and rapid adaptation [31]. In humans, functional groups of genes such as “Transcription Factor and/or Development” and “Receptor and/or Membrane” have been identified as enriched in STRs [32]. Similar enrichment in functional groups have been identified in yeast [33], fruit fly [34] and in transcription and translation in plants and algae [35]. However, the degree to which Atlantic cod and other species of the Gadiformes share the same genomic distribution of these STRs within functional groups as in human and other species, is currently unknown and will require high-quality genome assemblies of additional gadiform species.

In this study, we have generated a highly contiguous genome assembly for haddock (*Melanogrammus aeglefinus*) using a combination of PacBio and Illumina reads. Our aim was to perform a comparative genomic analysis with the only other currently available highly contiguous gadiform genome assembly – that of Atlantic cod. The haddock assembly is comparable to the Atlantic cod assembly with regards to contiguity and gene content. Using this new assembly, we have further investigated the immune gene repertoire and the impact of STRs in Gadiformes. We show that ray-finned fish – including cod and haddock – are enriched for genes with STRs in functional groups (based on Gene Ontology) such as transcription factors. In addition, the codfishes are significantly enriched for STRs in functional groups associated with signal transduction. Comparative analyses indicate a general expansion of the *NLR* genes in all teleosts, with possible lineage-specific expansions in zebrafish, stickleback and the codfishes.

## Results

### Assembly

Approximately 160x coverage of Illumina paired end reads and 20x coverage of PacBio reads were assembled with the Celera Assembler [36] resulting in a contig assembly (see Methods). All Illumina reads were mapped to the contig assembly with the Burrows-Wheeler Aligner (BWA) [37], and the scaffold module from String Graph Assembler (SGA) [38] was used to scaffold the contigs. To reduce gaps and to improve the accuracy of the consensus sequence, all Illumina reads were mapped to the scaffold assembly, and Pilon [39] was run to improve the contigs using high-coverage short-read information. Table 1 lists the statistics of the final assembly (also referred to as melAeg) and that of two assemblies from [26] for comparison. The melAeg assembly has shorter contigs and scaffolds than gadMor2, but approximately the same numbers of genes are found with CEGMA [40,41] and BUSCO [42]. The GM_CA454PB assembly was one of the four assemblies combined to make gadMor2 [26], and was created in a similar way to melAeg. It has similar contig and scaffold lengths, but fewer conserved genes were found by CEGMA and BUSCO.

**Table 1:** Genome assembly statistics for haddock (melAeg) compared with two assemblies of Atlantic cod, one draft based on PacBio and 454 reads (GM_CA454PB) and the final gadMor2 assembly

|  | melAeg | GM_CA454PB | gadMor2 |
| --- | --- | --- | --- |
| Length assembly (Mbp) | 653 | 681 | 644 |
| N50 scaffold (kbp) | 209 | 272 | 1,150 |
| N50 contig (kbp) | 78 | 95 | 116 |
| CEGMA complete (% of 458 genes) | 439 (96 %) | 431 (94 %) | 435 (95 %) |
| BUSCO single | 4,041 (88 %)<sup>1</sup> | 3,819 (83 %)<sup>1</sup> | 4,160 (91 %)<sup>1</sup> |
| BUSCO duplicated | 128 (2.8 %)<sup>1</sup> | 117 (2.6 %)<sup>1</sup> | 127 (2.8 %)<sup>1</sup> |
| BUSCO fragmented | 203 (4.4 %)<sup>1</sup> | 359 (7.8 %)<sup>1</sup> | 139 (3.0 %)<sup>1</sup> |
| BUSCO missing | 212 (4.6 %)<sup>1</sup> | 289 (6.3 %)<sup>1</sup> | 158 (3.4 %)<sup>1</sup> |

<sup>1</sup>% of 4584 genes

### Annotation and identifying orthologous genes

An iterative automatic annotation with MAKER [43,44] using an Illumina based transcriptome of haddock created from reads sequenced in [45], and proteins from UniProt/SwissProt [46], annotated 96,576 gene models. InterProScan [47] was run on the predicted proteins of these, and gene names were allocated based on match with proteins in UniProt/SwissProt. We created a filtered set where all genes had an Annotation Edit Distance (AED) [48] of less than 0.5 (where 0.0 indicates perfect accordance between the gene model and evidence (mRNA and/or protein alignments), and 1.0 no accordance). This resulted in 27,437 gene models.

We used OrthoFinder [49] to create a catalogue of orthologous genes, inferring them based on the predicted proteins of different species. We included the following species from Ensembl r81: Amazon molly (*Poecilia formosa*), cave fish (*Astyanax mexicanus*), Atlantic cod (*Gadus morhua*; gadMor1), fugu (*Takifugu rubripes*), medaka (*Oryzias latipes*), platyfish (*Xiphophorus maculatus*), spotted gar (*Lepisosteus oculatus*), stickleback (*Gasterosteus aculeatus*), tetraodon (*Tetraodon nigroviridis*), tilapia (*Oreochromis niloticus*) and zebrafish (*Danio rerio*), in addition to haddock and the most recent Atlantic cod genome assembly (gadMor2). For each gene, only the longest protein isoform was used. 281,838 proteins were placed into 17,519 orthogroups, with 20,661 proteins without a match. Cod and haddock have 11,500 groups in common (at least one protein from each species). See Supplementary Table 1 for the number of orthogroups shared between the other species-pairs.

### Genetic variation and historic effective population size

To be able to compare the heterozygosity rate between haddock and cod, we mapped the Illumina reads of the two species from [9] against the assemblies with BWA [37], and called SNPs (single nucleotide polymorphisms), MNPs (multi-nucleotide polymorphisms), indels (insertions and deletions) and complex regions (composite insertion and substitution events) with FreeBayes [50]. Haddock had 40 % more SNPs than cod (gadMor 2), with even larger differences in MNPs, indels and complex variants (Table 2).

**Table 2:** Number of variants called for the assemblies of haddock and cod. In parentheses the number of variants are given per bp, i.e. as nucleotide diversity.

|  | Haddock | Cod |
| --- | --- | --- |
| SNPs | 3,552,609 ($5.4 \times 10^{-3}$) | 2,506,699 ($3.9 \times 10^{-3}$) |
| MNPs | 127,929 ($0.2 \times 10^{-3}$) | 88,869 ($0.1 \times 10^{-3}$) |
| indels | 1,013,087 ($1.6 \times 10^{-3}$) | 608,828 ($0.9 \times 10^{-3}$) |
| complex | 300,678 ($0.5 \times 10^{-3}$) | 173,128 ($0.3 \times 10^{-3}$) |

While we have investigated only one individual per species, in general there is a correlation between nucleotide diversity of one individual and effective population size [51]. We used the Pairwise Sequentially Markovian Coalescent (PSMC) program [52] to infer the historic effective population size for the two species (Figure 1). We used a generation time of 10 years for cod and 6 years for haddock [53] with mutation rates derived from the phylogeny used in [9]. From this we found that haddock has an approximately 2.5 times larger historic effective population size than cod (Figure 1).

**Figure 1:**
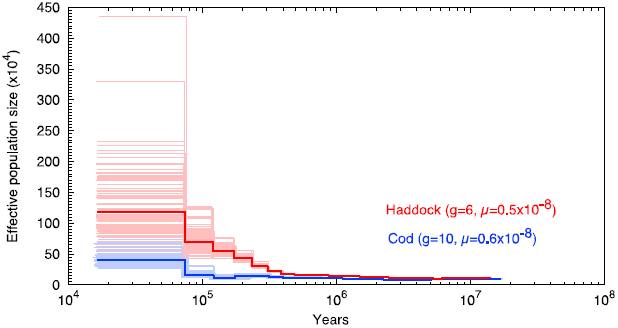
The historic effective population sizes in cod and haddock.

The analysis also includes the time before the two species split, as inferred by PSMC. Haddock is marked in red and cod in blue. Each analysis has been run with 100 bootstrap replicates, shown as pale versions of the main color. The time-span is ranging from approximately 20 million to 20,000 years ago.

### The *TLR* repertoire

Cod and haddock in general display the same TLR repertoire (Table 3). There is a difference of one or two gene copies for the cod assembly compared to what has been reported previously [13]. Our search criteria were quite strict, and the underlying assemblies were different (GM_CA454PB in [13], gadMor2 here), so some discrepancy can be expected.

**Table 3:** Number of full-length *TLR* genes found in the haddock and cod assemblies. Additional incomplete copies (≥60% of the entire gene) are indicated in parenthesis.

|  | Haddock | Cod |
| --- | --- | --- |
| TLR1/6 | 0 | 0 |
| TLR2 | 0 | 0 |
| TLR3 | 1 | 1 |
| TLR4 | 0 | 0 |
| TLR5 | 0 | 0 |
| TLR7 | 1(2) | 3 |
| TLR8 | 8(1) | 9 |
| TLR9 | 5(1) | 4(1) |
| TLR14 | 1 | 1 |
| TLR21 | 2 | 1(1) |
| TLR21beta | 0 | 0 |
| TLR22 | 5(1) | 10 |
| TLR23 | 1 | 2 |
| TLR25 | 4 | 5 |
| TLR26 | 0 | 0 |

Thirty-six full-length *TLRs* were identified for cod, whereas 28 were identified for haddock (Table 3). For both species, *TLRs* 1/6, 2, 4, 5, 21beta and 26 were not present. The gene numbers for most of the *TLRs* (*TLR* 3, 7, 9, 14, 21, 22, 23 and 25) were similar between both species. In contrast, cod had a significantly higher number of *TLR22* (10) than haddock (5).

### The *MHCI* repertoire

The number of *MHCI* loci has previously been characterized in cod, using both qPCR and read-depth comparisons, and 80-100 and ~70 copies were estimated, respectively [9,10]. By using read-depth comparisons for haddock, ~30 copies were calculated for this species [9]. Only two copies of *MHCI* were found in the first version of the cod genome assembly (gadMor1) [10]. We used the new assemblies of cod and haddock to investigate the number of copies of *MHCI.*

We inferred the presence of *MHCI* based on the occurrence of the three alpha domains of MHCI, including the most conserved alpha-3 domain. We found 13 regions with all three exons in cod, and 10 such regions in haddock. One significant difference between the two species was the number of occurrences of isolated alpha domains, suggesting potentially more copies of *MHCI* in cod (Table 4). Because these genes occur in multiple copies within the genome, the genome assembler might consider them as repeats [21], potentially resulting in fragmented assembly of these genes. We found up to 20 copies of *MHCI* (sum of all hits) in haddock, and 53 in cod, i.e., 66 % and 76 % of the previous estimated number of *MHCI* copies in haddock and cod, respectively [9].

**Table 4:**
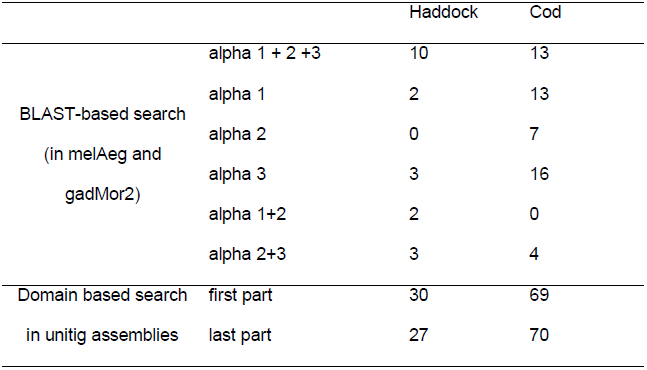
The number of *MHCI* found in the haddock and cod assemblies based on different criteria. The BLAST-based reports open reading frames for the hits in the final assemblies, while the domain based report the number of domains found in the unitig assemblies that underlie the final assemblies.

Celera Assembler, the assembler used for assembling melAeg and GM_CA454PB, outputs so-called unitigs in addition to outputting contigs and scaffolds. Unitigs are sequences that are either unique in the genome or are collapsed repeated sequence. These are incorporated into contigs based on different rules (e.g., likelihood of being a repeat). Often, the contigs only contain a subset of the unitigs, and therefore could contain fewer genes. We translated the unitigs assemblies of melAeg and GM_CA454PB into all six reading frames with transeq [54] and searched these with the MHCI PFAM [55] domain PF00129 using HMMER [56]. For cod and haddock, the domain spans two exons, thus we counted occurrences of the first and last part of the profile found in the assemblies (Table 4). We found 27 copies of the first part of the domain and 30 copies of the last part in haddock and 69 and 70, respectively, in cod, approximately the same as in [9]. It is likely that some of these are collapsed because of the repeated nature of *MHCI* genes.

### Expansion of *NLRs* in teleosts

The zebrafish has a lineage-specific expansion of the *NLRs* [57], but it is unclear how many copies are found in other teleost genome assemblies. We investigated the *NLRs* with several approaches. First, we ran InterProScan [47] on the longest protein per gene to annotate protein domains. We parsed the output and counted occurrences of the PFAM [55] domains PF05729 (NACHT domain) and PF14484 (Fish-specific NACHT associated domain, FISNA) (Figure 2). Second, we translated the assemblies into all six reading frames with transeq [54] and used these to search for the NACHT and FISNA domains using HMMER [56]. For all species, the number of domains identified was substantially elevated when scrutinizing the assemblies compared to the predicted proteins (Figure 2). For example, in platyfish the number of NACHT domains increased from 29 to 120. The reported numbers show a large variation in copy number between the different species (Figure 2), with large difference between relatively closely related species, such as tetraodon and fugu, or cod and haddock, where there are three times as many copies in cod compared to haddock.

**Figure 2:**
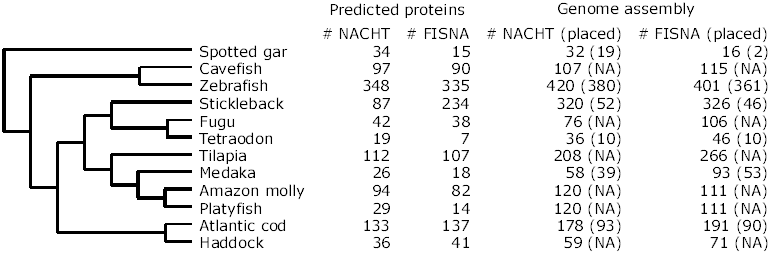
NACHT and FISNA domains content in predicted proteins and genome assemblies for the different species.

HMMER hits had to be >75 % of the length of the domain to be reported here. Some species have scaffolds ordered and organized into chromosomes/linkage groups, i.e., placed. For these species the number of domains found in placed scaffolds are also reported. NA: Not applicable.

For the species with contigs/scaffolds placed into either linkage groups or chromosomes (cod, stickleback, zebrafish, spotted gar, medaka and tetraodon) we counted the number of genes where the relevant domains were found in either placed (i.e. in the linkage map) or unplaced sequences (Figure 2, Supplementary Tables 5-10). We found that many of the sequences with these kinds of domains are unplaced, as previously reported [18,57]. While zebrafish has a majority of domains in placed sequences, most sequences in stickleback with FISNA and NACHT domains are not placed. About half the sequences are placed in cod, while most sequences are placed in the other species.

**Table 5:** The number of hits for NACHT and FISNA domains in the unitig assemblies for cod and haddock. Substantially more hits are found in the unitigs that in the contigs of the final assemblies, indicating that many of the unitigs are not included, possibly because they are categorized as repetitive sequence.

|  |  | Haddock | Cod |
| --- | --- | --- | --- |
| NACHT | all | 613 | 656 |
|  | >50 % domain length | 224 | 264 |
|  | >75 % domain length | 121 | 140 |
| FISNA | all | 611 | 552 |
|  | >50 % domain length | 553 | 505 |
|  | >75 % domain length | 384 | 359 |

There are multiple reasons for a genome to not assemble properly, but repeated sequence is one of the most influential [21]. Genes occurring in multiple copies such as *NLRs* are indistinguishable from any other repeated sequence for the assembler. One consequence of this is that some of these unplaced contigs/scaffolds would have higher coverage in reads than average since they basically are collapsed repeats. For haddock and cod we have sequencing read data available, and we estimated and plotted the average coverage for all sequences with the FISNA domain (Figure 3). Many of the sequences shorter than 100,000 bp show a higher than average coverage. This is especially the case for those sequences around 10,000 bp, and indicates that these contain multiple copies of the FISNA domain, i.e. these contain collapsed copies.

**Figure 3:**
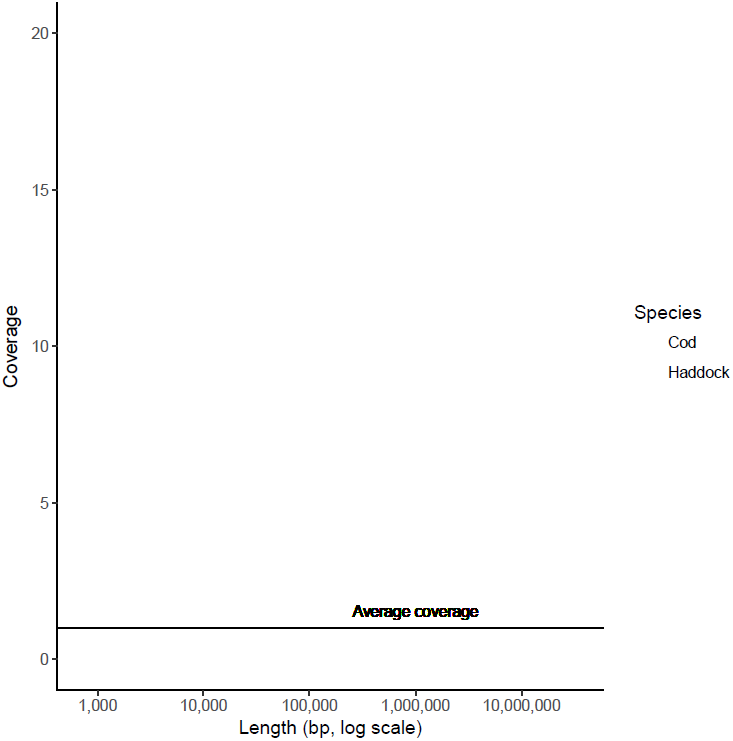
Relationship between length and coverage of reads for sequences harboring the FISNA domain.

Coverage has been normalized for each species by dividing the coverage for each sequence with the average for that species. The average lengths of genes with the FISNA domain is 17 kbp in cod and 14 kbp in haddock, and the increased coverage in sequences about this length might indicate that there are multiple, very similar regions with these genes in the two species. The cod sequences larger than 10 Mbp represent the linkage groups. Cod is plotted with red and haddock in blue. The x-axis is log(10)- transformed since the sequences span from 700 bp to more than 20 Mbp.

Due to differences in the assembly strategy, the haddock assembly contains fewer short contigs than the cod assembly (Supplementary Note 1). We investigated the unitig assemblies for cod and haddock with the NACHT and FISNA domains, with the same approach as used for *MHCI* for unitig assemblies (Table 5). This approach reports around 600 copies of each of the domains in both species. The NACHT domain is longer (166 aa) than the FISNA domain (72 aa), and while the total number of hits is similar between the two domains, there are significantly fewer NACHT domains found at >75 % of the domain length. The short hits for the NACHT domain are predominantly found on unitigs shorter than 500 bp, suggesting that these are collapsed.

### Investigating the STR content of the haddock genome assembly

We investigated the amount of short tandem repeats (STRs) in the haddock genome assembly, compared to cod and other ray-finned fishes. We used Phobos [58] to annotate all STRs with an unit size of 1-10 bp. Haddock has an even higher density of STRs in its genome assembly compared to cod, 96,364 bp/Mbp in haddock and 80,706 bp/Mbp in cod (Figure 4A). The amino acid coding parts of the genome also contain a high proportion of STRs, 25,639 bp/Mbp in haddock and 16,501 bp/Mbp in cod. This mostly consists of dinucleotide repeats, but both cod and haddock have approximately 6,000 bp/Mbp of trinucleotide STRs in protein coding regions, compared to 530 bp/Mbp in medaka, and up to 934 bp/Mbp in zebrafish with the other fishes harboring intermediate amounts (Figure 4B). Cod and haddock also have higher frequencies (loci/Mbp) of STRs in the assemblies (Figure 4C and Supplementary Table 2), and in the protein coding regions (Figure 4D). By using the overlap between annotated STRs and genes, we also report the number of genes with one or more STR for these species (Supplementary Table 3).

**Figure 4:**
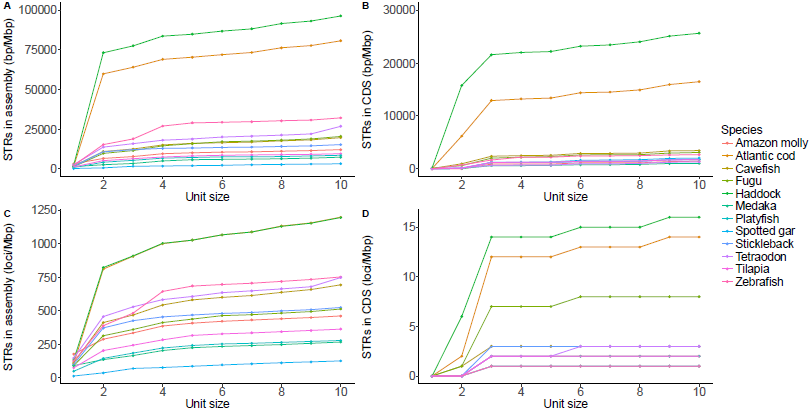
Cumulative plot of the density (bp/Mbp) and frequency (loci/Mbp) of short tandem repeats (STRs).Shown is the STR content per unit size in the whole assembly and CDS for different teleosts. Most of the STR contents in the whole assembly in cod and haddock are dinucleotide repeats, but there are about equal amounts of dinucleotide and trinucleotide repeats in coding sequence. A. Density of STRs in the genome assembly (bp/Mbp). B. Density of STRs in protein coding regions (bp/Mbp). C. Frequency of STRs in the genome assembly (loci/Mbp). D. Frequency of STRs in the in protein coding regions (loci/Mbp).

For haddock and cod, we were also able to find indels (called by FreeBayes) and STRs in protein coding regions, and where these structural variants overlap. We found STRs of all unit sizes in the protein coding regions (Figure 4D), but those STRs with unit sizes that do not create frame shifts, such as tri-, hexa- and enneanucleotides, are most interesting from a functional perspective. Of these, the vast majority are trinucleotides, and we restricted our analysis to these. We found 581 genes with an indel of size 3 in a trinucleotide repeat in haddock (2.1 %) and 660 genes in cod (2.9 %), i.e. these are heterozygous in these two individuals.

### Between-species comparisons of STR enrichment in genes

Cod and haddock have a much larger proportion of their protein coding sequence in dinucleotide and trinucleotide STRs compared to other species (Figure 4). In the process of annotating a genome, many genes are assigned a gene ontology term (GO term), describing the processes the protein encoded by that gene is involved in. We wanted to investigate if genes with STRs are randomly spread across different GO groups, or if some GO groups in some species are enriched for genes with STRs. Fisher’s exact test was used to perform pairwise comparisons of the number of genes with STRs and the number of genes without STRs between each species (Figure 5 for examples, Supplementary Figure 1 and Supplementary Table 4 for details). Of the 2,748 GO terms in the dataset, there are significant differences between species in 74 GO groups after correcting for multiple testing (false discovery rate with Benjamini/Yekutieli). For many of these, haddock and cod differ significantly from all other species, but not from each other (Supplementary Table 4). These include protein kinase activity (GO:0004672), G-protein coupled receptor activity (GO:0004930), signal transduction (GO:0007165), metabolic process (GO:0008152) and transmembrane transport (GO:0055085).

**Figure 5:**
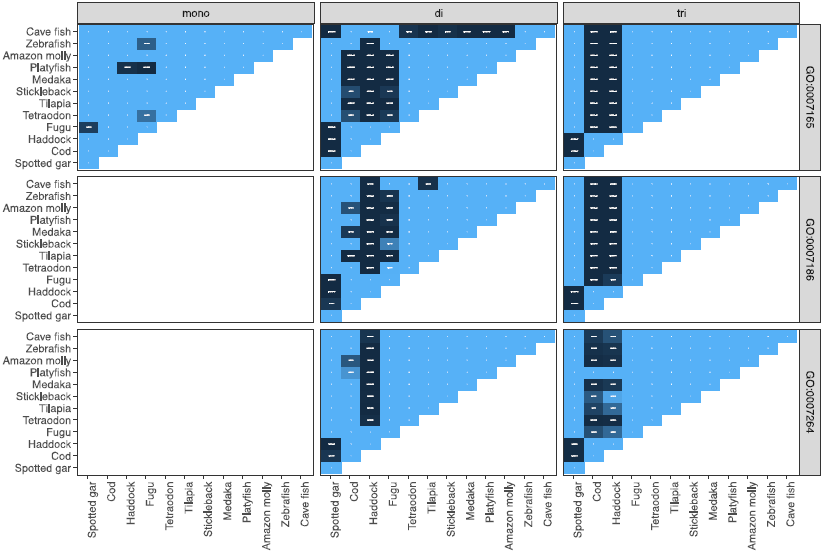
Pairwise Fisher’s exact test for some gene ontology groups and for some unit sizes. See supplementary Figure 1 for the entire figure with 74 GO groups and unit sizes 1-10 bp, and Supplementary Table 4 for the GO groups where haddock and cod differ significantly from the other species. Shown here are GO:0007165 (signal transduction), GO:0007186 (G-protein coupled receptor signaling pathway) and GO:0007264 (small GTPase mediated signal transduction) for tandem repeats in 1-3 bp unit sizes. In the white and blue areas there are no significant differences, but in the dark blue areas there are significant differences between two species. For GO:0007165 and GO:0007186 there is a significant difference (P<0.05) between cod and haddock and the other species, but not between cod and haddock, nor between cod and cave fish. For GO:0007264, this pattern is less apparent.

### Within species comparisons of STR enrichment within genes

To investigate enrichment and purification (under-representation) of STRs in GO terms, we used goatools [59] (Figure 6, Supplementary Figure 2). We corrected for multiple testing. For some terms, both cod and haddock are enriched, whereas this is not the case in the other species. These are cation channel activity (GO:0005261), regulation of signal transduction (GO:0009966), regulation of cell communication (GO:0010646), regulation of signaling (GO:0023051), regulation of Rho protein signal transduction (GO:0035023), regulation of Ras protein signal transduction (GO:0046578), regulation of response to stimulus (GO:0048583), regulation of small GTPase mediated signal transduction (GO:0051056), regulation of intracellular signal transduction (GO:1902531). These are mainly in the hierarchy above regulation of Rho protein signal transduction (GO:0035023), as well as cation channel activity (GO:0005261).

**Figure 6:**
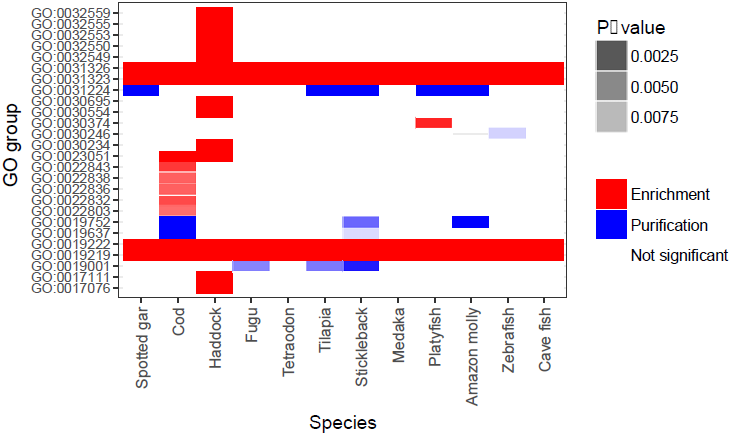
An example of gene ontology terms significantly enriched for genes with trinucleotide tandem repeats in different species.

Trinucleotide tandem repeats are repeats that can vary in number of repeat units without causing frameshifts in the protein. Only tests with P<0.01 are colored. Red signifies enrichment, i.e. more trinucleotide repeats than expected, and blue purification, i.e. less than expected. See Supplementary Figure 2 for the complete analysis.

## Discussion

### A highly contiguous genome assembly for haddock

Here we have taken advantage of long and short read technologies to produce an annotated and highly contiguous assembly of the haddock genome, with comparable gene content and assembly statistics to the recently released Atlantic cod genome assembly [26] (Table 1). The genic completeness of the assembly is high, as seen by the BUSCO score, where >90 % of the 4,584 genes are found complete (Table 1). PacBio reads span more repeated regions than Illumina reads, and the contig N50 is therefore longer for the haddock assembly than other fishes sequenced with only Illumina reads, for instance the Asian arowana [60] and the seahorse [5]. With the increased affordability, availability and usage of such long-read sequencing technologies as PacBio [61] and Oxford Nanopore [62] reads, more complete assemblies for diverse species are likely to arrive in near future.

### Increased number of tandem repeats in codfishes

Several studies have shown the Atlantic cod genome has a high STR content [63-65]. The first version of the cod genome assembly [10] was fragmented, and STRs have recently been identified as the main factor causing this fragmentation [26]. Since STRs have a high mutation rate, their presence in genes might disrupt normal gene product function, as seen for the multitude of human diseases due to large expansions in STRs [66]. Surprisingly, while both cod and haddock have a high density and frequency of STRs in the assembly overall, they also have a substantial amount of STRs in protein coding regions compared to other ray-finned fish (Figure 4). STRs shrink and expand by DNA polymerase slippage or recombination [27], but a repeated motif has to be present for this to happen. A short tandem repeat might be created by a mutation (changing ATAG to ATAT), or as the result of transposable element activity [67]. Further work is needed to investigate the basis for the high STR content in codfishes.

STRs are present in almost twice as many genes in cod and haddock compared to the other ray-finned fishes (Supplementary Table 3). Specifically, in around 8,000 genes in codfishes compared to 1,500- 4,000 in the other species. This is almost twice as many as in humans (4,500) [29]. In humans, genes connected to processes such as transcriptional regulation, chromatin remodeling, morphogenesis, and neurogenesis have been found enriched for STRs [32,68]. Similar enrichment has been found in other species, such as yeast [33], fruit fly [34] and plants and algae [35]. In the fish species investigated here, there is enrichment in genes with STRs in functional (Gene Ontology) groups primarily concerned with transcription, similar to previous studies [33-35] (Supplementary Figure 2). One example is the transcriptional regulator Ssn6 in yeast, where increased length of a polyglutamine tract (encoded by a STR), was positively correlated with increased expression of some target genes, and negative correlated with others [69]. Haddock and cod have significantly larger proportions of genes with STRs in GO groups associated with genes encoding proteins involved in signal transduction compared to the other species. These GO groups contain a higher proportion of genes with STRs than expected with comparing GO groups per species. This is also true when comparing GO groups between species. Many of these functional groups are connected to small GTP-binding proteins such as regulation of Rho protein signal transduction (GO:0035023), regulation of Ras protein signal transduction (GO:0046578), and regulation of small GTPase mediated signal transduction (GO:0051056). The small GTP-binding proteins are involved in regulation of processes such as gene expression, cytoskeletal reorganization, intracellular vesicle trafficking and cytokinesis [70,71]. The regulation of the activity of small GTPases are mainly performed by GTPase-activating proteins (GAPs) and guanine nucleotide-exchange factors (GEFs) by suppression (GAPs) or promotion (GEFs) of the GTPase’ activity [72]. For instance, in humans, 81 GEFs and 67 GAPs [73] regulate the activity of the 22 Rho GTPases [74]. Some of the small GTPases are important for proper immune function [75,76], by regulating chemotaxis and phagocytosis [77]. In mammals, the GTPase RhoA is important for TLR signaling, specifically for TLR2 and TLR4 [77]. Thus, between two populations of codfishes, adapted to different environments, there may potentially be variation in immune responses based on length variations of STRs in GEFs and GAPs.

### Historic effective population size and STRs

Many marine fish with a pelagic life style are characterized by large effective population sizes [78]. Atlantic herring has an estimated effective population size of approximately 1 million and a nucleotide diversity of 0.32 % [4], similar to cod with effective population size around 400,000 and 0.39 % nucleotide diversity and haddock at around 1.1 million and 0.54 % nucleotide diversity (Table 1). Intriguingly, herring seems to have a high amount of STRs (Supplementary File E in [4]), suggesting that the life history strategies of cod, haddock and herring might facilitate a high density and frequency of STRs. The high effective population sizes in these species would imply low genetic drift and more efficient selection.

With around 760,000 STR loci in haddock and cod (Supplementary Table 2), the majority are likely to be highly polymorphic in such large haddock and cod populations. In a study of over 1,000 human individuals, most of the 700,000 STR loci sequenced were polymorphic [29], although constraints were apparent for mutations in coding sequences [28]. Haddock and cod (Figure 1) have at least ten times the historic effective population size of humans [52], and their high fecundity would generate many STR variants for each generation. We find trinucleotide indels in STRs in 2-3 % of the genes, i.e., they have different length variants of the STRs in these genes. With such large effective populations and few barriers, genetic drift is weak, and local populations should respond to even weak selection [78]. There are studies suggesting STR loci are under selection in cod [79,80]. Most tools for genome-wide investigations of selection have focused on SNPs, but methods for selection on STRs have been developed [81]. With high accuracy STR genotyping [82,83] and resequencing data from different populations or controlled experiments over several generations, we suspect substantial numbers of STRs under selection will be found.

### The MHCI and TLR repertoire in haddock and cod

In the first cod genome assembly, only two *MHCI* classical U-lineage genes were found, despite qPCR indicating around 100 copies [10]. Other investigations have also estimated a large number of *MHCI* copies in cod [9,84,85], but these have either investigated transcriptional data or read depth comparisons between *MHCI* loci and single-copy genes. [9] estimated around 30 copies in haddock and 70 in cod. We found similar numbers to those predicted by [9] using our unitig assemblies of the same species; however in contrast a much lower number was found in the final assemblies. In the cod assembly, seven of the in total thirteen *MHCI* copies with complete alpha domains are located on unplaced contigs/scaffold in the gadMor2 assembly (data not shown). Their numbers are likely to be underestimated because the unplaced contigs/scaffold often have a higher read depth, indicating that these contain multiple, collapsed copies. Using PacBio reads in both the haddock and the cod assemblies likely substantially contributed to the more complete representation of *MHCI* genes, compared to the previous cod genome assembly. The Asian seabass, another assembly based on PacBio reads, resulted in “a more continuous cluster of MHC-class I genes compared to the well-assembled *G*. *aculeatus* [three-spined stickleback] genome” [24], highlighting the importance of long reads for properly capturing these regions of the genome. In contrast, the *TLR* repertoire is by and large similar between haddock and cod. The only main difference is TLR 22; with twice as many copies in cod (10 vs. 5). We were unable to perform the domain-based search for *TLRs*, since they do not have a *TLR*-specific domain. The TIR domain (PFAM domain PF01582), the most likely candidate, is also found in the large interleukin-1 receptor family [86].

### The high copy number of *NLRs* in teleosts and genome assembly

In this study we enumerate genes (putative *NLRs*) with the NACHT (PFAM domain PF05729) and FISNA (PF14484) domains. These two domains together characterize a family of proteins substantially expanded in zebrafish with around 400 copies [57] and indications of substantial expansions in other teleosts as well [18,87,88].

For genome assemblers, identical or highly similar sequences occurring in multiple locations in a genome are indistinguishable from repeated sequence such as for example transposable elements. Depending on the sequencing strategy and assembler, these may introduce gaps into an assembly because the assembler is unable to place them correctly and they might be collapsed as a single contig/scaffold [21]. In general, genome assemblers might treat the large amount of *NLR* genes in these species as repeated sequence, and thus be unable to place them into scaffolds. For the species with genome assemblies in linkage groups or chromosomes, we looked at the contigs/scaffolds that were placed into these versus those that were not (Figure 2). Even with the large number of genes (>400), only 10 % of the putative *NLRs* are unplaced for zebrafish. This is likely due to its sequencing and assembly strategy, with tiling of individually sequenced and assembled bacterial artificial chromosome clones [90]. For cod, about 50 % the contigs/scaffolds with putative *NLRs* are unplaced, and for stickleback about 15 % are unplaced. The stickleback genome assembly is based on 9x coverage with Sanger sequencing reads [91], which may result in a more fragmented assembly than using PacBio reads (as for cod) or clones (zebrafish) because Sanger sequencing reads are shorter.

The numbers of putative *NLRs* from Figure 2 should be interpreted with caution. It is likely that all species have some or several of the gene copies collapsed [18]. For cod and haddock, we mapped reads back to the assembly, and investigated the coverage for all sequences (Figure 3). There are many contigs/scaffolds with more than 5 times coverage compared to the average in the assemblies, and the numbers of putative *NLRs* are likely underestimated. Even though these two assemblies are highly contiguous and have been created with the use of PacBio reads, multi-copy genes such as *NLRs* may still be problematic. We also investigated the content of the unitig assemblies for cod and haddock, and found similar numbers of *NLRs* between the two species (Table 5). The difference between the unitig assemblies and the final assemblies are because of differences in assembly processes (Supplementary Note 1), where the final haddock assembly contains fewer short contigs. Most likely the *NLR* content of the two codfishes is highly similar. The numbers of *NLRs* are likely severely underestimated in most currently investigated ray-finned fish. Assemblies of higher quality are needed to properly investigate this intriguing family of innate immune genes.

It is unclear how such large gene families as the *NLRs* in zebrafish evolved [92]. In zebrafish, the majority of *NLRs* are located on one chromosome 4 arm [57] (Supplementary Table 6). Although the other assemblies are of lower quality than the zebrafish genome, there are no clear patterns of chromosomal enrichment in *NLRs* in other ray-finned fishes. Possible exceptions are medaka with 33 FISNA domains found on linkage group 2 (Supplementary Table 9) and stickleback with 12 FISNA and NACHT domains found on groupXIII (Supplementary Table 7). For Atlantic cod, the *NLRs* are evenly divided across linkage groups (Supplementary Table 5). Further, tetraodon (Supplementary Table 10) and spotted gar (Supplementary Table 8) have relatively few copies in total.

## Conclusions

Our study provides new insight into elements of genomic architecture in codfishes. The haddock genome contains an even higher density of STRs than the Atlantic cod genome. Further, certain classes of genes are enriched for STRs in both Atlantic cod and haddock, but not in the other published fish genome assemblies. With the large effective population sizes of cod and haddock, these STRs are likely polymorphic and represent a large reservoir of genetic variation. Additionally, for copy number estimations of highly expanded genes, such as the *NLR* genes, we discovered that the genome assemblies of most teleosts do not accurately represent these. Thus, the expanded nature of such gene families most likely confound genome assemblers, at least when based on Illumina reads or moderate coverage of PacBio reads. However, investigation of unitig assemblies of cod and haddock shows substantial higher copy numbers than the final assemblies. Most likely, the teleost genome assemblies available represent severe underestimations of the number of NLR genes. Better genome assemblies, i.e. created with sufficient long read coverage in combination with linked reads [93], optical mapping [61,94] and/or chromosome conformation [23], should facilitate proper characterization of the *NLR* content as well as other teleost multicopy genes, unraveling their evolutionary past.

## Materials and methods

### Sampling and sequencing

The sequenced individual, a wild caught specimen approx. 1.3 kg belonging to the North-East Artic haddock population, was sampled near the Lofoten Islands (N68.04 E13.41), outside of its spawning season (in July 2009). The fish were humanely euthanized before sampling in accordance with the guidelines set by the‘Norwegian consensus platform for replacement, reduction and refinement of animal experiments’ (<u>www.norecopa.no</u>). The DNA was extracted from spleen (stored on RNALater) using a standard high salt DNA extraction protocol.

200 bp insert size paired end libraries were constructed with Illumina DNA paired end sample preparation reagents and sequenced at the McGill University and Génome Québec Innovation Centre, both 100 bp and 150 bp reads. The 3 kbp and 10 kbp insert size libraries were prepared with the Illumina Mate Pair gDNA reagents and sequenced at the McGill University and Génome Québec Innovation Centre with 100 bp reads. All Illumina libraries were sequenced on the HiSeq 2000 using V3 chemistry.

PacBio SMRT sequencing was performed on a PacBio RS II instrument (Pacific Biosciences of California Inc., Menlo Park, CA, USA) at the Norwegian Sequencing Centre (NSC, www.sequencing.uio.no/). Long insert SMRTbell template libraries were prepared at NSC according to PacBio protocols. In total, 24 SMRT-cells were sequenced using P6v2 polymerase binding and C4 sequencing kits with 120 min acquisition. Approximately 16.4 Gbp of library bases were produced.

## Assembly

### Genome assembly

Meryl from Celera Assembler 8.3rc2 [36] was used to count k-mers in the paired end Illumina libraries. All Illumina paired end reads were sequenced from the same DNA library, with insert size around 200 bp. Because of this overlapping reads were merged with FLASH v1.2.3 [95].

The merTrim program [26], also from Celera Assembler, was used to correct the output from FLASH, the merged and unmerged Illumina reads. The raw, uncorrected PacBio whole genome shotgun reads were separately trimmed by the overlap-based-trimming module in Celera Assembler [36]. The trimmed Illumina and PacBio reads were assembled together with Celera Assembler resulting in a contig assembly, following [26]. All Illumina reads were mapped to the contig assembly using BWA mem v0.7.9a [37], and the scaffold module from SGA (github snapshot June25th_2014) [38] was used to scaffold the contigs. All Illumina reads were again mapped to the scaffold assembly, and Pilon v1.16 [39] was applied, reducing some gaps and recalling consensus.

### Transcriptome assembly

All RNA-seq data from [45] (Sequence Read Archive at NCBI with Accession ID: PRJNA328092) was assembled with Trinity v2.0.6 [96].

### Validation of genome assembly

CEGMA v2.4.010312 [40,41] and BUSCO v2 [42] with an actinopterygii specific gene set were run on the genome assembly to asses the amount of conserved eukaryotic genes.

## Annotation

### Repeat library

A library of repeated elements was created as described in [26]. RepeatModeler v1.0.8, LTRharvest [97] part of genometools v1.5.7 and TransposonPSI were used in combination to create a set of putative repeats. Elements with only a match against an UniProtKB/SwissProt database and not against the database of known repeated elements included in RepeatMasker were removed. The remaining elements were classified and combined with known repeat elements from RepBase v20150807 [98].

### Annotation

Three different *ab initio* gene predictors were trained. GeneMark-ES [99] v2.3e on the genome assembly, SNAP v20131129 [100] on the genes found by CEGMA, and AUGUSTUS v3.2.2 [101,102] on the genes found by BUSCO. MAKER v2.31.8 [43,44] used the trained gene predictors, the Trinity transcriptome assembly, the repeat library and proteins from UniProtKB/SwissProt r2016_3 [46] for a first pass [103] annotation of the genome assembly. The result of the first pass was used to retrain SNAP and AUGUSTUS, and a second iteration was performed using the same set-up.

The protein sequences from final output of MAKER was BLASTed against the UniProtKB/SwissProt proteins and InterProScan v5.4-47 [47] was used to classify protein domains in the protein sequences. Finally, the output of MAKER was filtered on AED, keeping only genes/proteins with an AED less than 0.5 (where 0.0 indicates perfect accordance between the gene model and evidence (mRNA and/or protein alignments), and 1.0 no accordance).

## Finding orthologues

We downloaded all genome assemblies, cDNA and protein fasta files for all fishes at Ensembl release 81 (Amazon molly, cavefish, Atlantic cod (gadMor1), fugu, medaka, platyfish, spotted gar, stickleback, tetraodon, tilapia and zebrafish), and extracted the longest protein using a custom script (get_only_longest_protein_per_gene.py) because some annotations provide multiple proteins per gene. We did an all-against-all BLASTP of the protein sequences of all the Ensembl fishes in addition to the new cod and haddock annotated proteins, following the default options as set by OrthoFinder. The results of this were used as input to OrthoFinder v1.0.6 [49].

## Investigating variants in the haddock and cod assemblies

Both haddock and cod were sequenced in the [9] study, and these 150 bp reads were mapped to the respective assemblies using BWA MEM v0.7.12 [37], and sorted using samtools v0.1.19 [104]. Bamtools v2.3.0 and the script ‘coverage_to_regions.py’ from FreeBayes v0.9.14 [50] were used to split the assembly into regions, and FreeBayes was run in parallel. Vcflib from a GitHub snapshot at 20140325 was used to filter the variants, and only variants with more than 20 in quality and 5 in depth were retained.

## Estimating historic effective population size

A GitHub snapshot from August25th 2015 of PSMC [52] was used together with samtools v1.1 and bcftools v1.2 on the mapped reads, and historic effective population size was inferred for cod and haddock. The mutation rates were estimated along the branches of the phylogeny reported in [9] and the generation times were set to 10 years for cod and 6 years for haddock [53].

## Identification of Toll-like receptors

Toll-like receptors (TLRs) are a key component of the innate immune response. The toll interleukine receptor (TIR) is the most conserved domain of the TLRs [105]. To determine candidate regions likely containing TLR genes, we aligned all TIRs protein sequences available on Ensembl and GenBank against the haddock and cod genome assemblies using TBLASTN from the BLAST+ suite [106] with an e-value cutoff of 1e-10. We then extracted 10,000 bp around the regions containing TIR like motifs. We used BLASTN to align coding sequences representative of all the TLRs classes against the candidate regions containing TLR copies. Here we report full-length TLR copies as well as partial copies (≥60% of the coding sequence).

## Identification of *MHCI*

We used the alpha-3 domain of the MHC I complex to identify the candidate regions containing *MHCI* genes in both haddock and Atlantic cod. We used TBLASTN to align alpha-3 coding sequences from Atlantic cod and zebra fish (*Danio rerio*) against the haddock and Atlantic cod genome assemblies, with an e-value threshold of 1e-10. We then extracted the region located 10,000 bp around the putative alpha-3 domains. We used BLASTN to align the extracted regions against the non-redundant nucleotide database on NCBI. Regions containing the three alpha domains of MHCI (α1, α2 and α3) were used as a proxy to determine the number of MHCI gene copy number.

To better assess the differences between the unitig assemblies and the final assemblies, we translated the unitigs assemblies of melAeg and GM_CA454PB (both are basis for the final assemblies) into all six reading frames with transeq from Emboss v6.5.7 [54], and used the PFAM v31.0 [55] domain PF00129 (Class I Histocompatibility antigen, domains alpha 1 and 2; MHCI) in HMMER v3.1b2 [56] to search the unitig assemblies for putative *MHCI* genes.

## Identification of *NLRs*

We ran InterProScan v5.4-47 [47] on the longest protein per gene to annotate protein domains. The default Ensembl annotation of these seemed out of date for several species, and with this procedure we had a more uniform dataset. We counted the occurrences of the PFAM v31.0 [55] domains PF05729 (NACHT domain) and PF14484 (Fish-specific NACHT associated domain, FISNA). In addition we translated the assemblies of all species into all six reading frames with transeq from Emboss v6.5.7 [54], and searched these with the NACHT and FISNA domains with HMMER v3.1b2 [56]. The species relationship in Figure 2 is derived from [9] and we used ETE3 [107] to plot the dendogram.

We used v1.3.1 of samtools [104] with the‘depth –a –a’ option to calculate the per base pair coverage of the assemblies, and used awk to calculate average depth per sequence and average for the whole assembly. We extracted all sequences with FISNA domains, and plotted length versus depth for these using ggplot2 [108] in the R environment.

As for *MHCI*, we searched the unitig assemblies of cod and haddock with the FISNA and NACHT domains.

## STRs in the assemblies and coding regions

We used Phobos v3.3.12 [58] to detect all TRs with unit size 1-10 bp in the assemblies. The output was in Phobos native format that was processed with the sat-stat v1.3.12 program, yielding files with different statistics and a gff file. The other settings were as used in [26].

We counted the number of different STRs in genes and number of genes with STRs by using bedtools[109] and overlaps between STRs and genes. For cod and haddock, we also counted the number of overlaps between trinucleotide TRs, indels of size 3 and genes.

## Enrichment of STRs in genes

For each gene ontology group we performed pairwise comparisons of the number of genes with STRs and total number of genes between the different species using Fisher’s exact test (implemented in SciPy [110]). We corrected for multiple testing using the Benjamini-Yekutieli [111] procedure of False Discovery Rate as implemented in statsmodels (http://www.statsmodels.org/stable/index.html). Of 2,748 gene ontology terms, we found significant differences in 74.

For each gene ontology group we also tested the enrichment or purification of STRs compared to amount of STRs all the genes in a species using goatools, and correcting for multiple testing with Benjamini-Yekutieli procedure of False Discovery Rate [59].

### Declarations

#### Ethics approval and consent to participate

We have adhered to all local, national and international regulations and conventions, and we respected normal scientific ethical practices.

#### Consent for publication

Not applicable.

#### Availability of data and materials

The genome assembly and annotation are available from FigShare: <u>https://doi.org/10.6084/m9.figshare.5182861</u>

Illumina sequencing reads are available from ENA at http://www.ebi.ac.uk/ena/data/view/PRJEB21701.

#### Authors’ contributions

OKT created the genome assembly and annotated it, performed all analyses involving STRs, the analyses using protein domains and wrote the first draft of the manuscript. MSOB performed the BLAST- based analyses with assistance of MHS. MHS, RBE and TF performed preliminary analysis of NOD-likreceptor genes. ES sampled and extracted RNA for RNA-seq. OKT, AJN and SJ designed the sequencing strategy. KSJ and SJ oversaw the project. SM, RBE and SJ conceived the study. All authors read and approved the final manuscript.

## Acknowledgements

All computational work was performed on the Abel Supercomputing Cluster (Norwegian metacenter for High Performance Computing (NOTUR) and the University of Oslo) operated by the Research Computing Services group at USIT, the University of Oslo IT-department (http://www.hpc.uio.no/). Sequencing library creation and high throughput sequencing was carried out at the Norwegian Sequencing Centre (NSC), University of Oslo, Norway. We especially thank Marianne H. S. Hansen for DNA extraction and Ave Tooming-Klunderud for PacBio RS II library preparation and sequencing, both affiliated NSC, University of Oslo. We also thank Mark Ravinet for critical reading of the manuscript.

